# Quinoxaline-Based Anti-Schistosomal Compounds Have Potent Anti-Malarial Activity

**DOI:** 10.1101/2024.04.23.590861

**Authors:** Mukul Rawat, Gilda Padalino, Tomas Yeo, Andrea Brancale, David A. Fidock, Karl F. Hoffmann, Marcus C. S. Lee

**Affiliations:** Biological Chemistry and Drug Discovery, Wellcome Centre for Anti-Infectives Research, University of Dundee, Dundee, United Kingdom; Wellcome Sanger Institute, Wellcome Genome Campus, Hinxton, United Kingdom; Department of Life Sciences (DLS), Aberystwyth University, Aberystwyth, United Kingdom; Swansea University Medical School, Swansea, United Kingdom; Department of Microbiology and Immunology, Columbia University Irving Medical Center, New York, New York, United States; Center for Malaria Therapeutics and Antimicrobial Resistance, Division of Infectious Diseases, Department of Medicine, Columbia University Irving Medical Center, New York, New York, United States; Department of Organic Chemistry, UCT Prague, Prague, Czech Republic

## Abstract

The human pathogens *Plasmodium* and *Schistosoma* are each responsible for over 200 million infections annually, being particularly problematic in low- and middle-income countries. There is a pressing need for new drug targets for these diseases, driven by emergence of drug-resistance in *Plasmodium* and the overall dearth of new drug targets for *Schistosoma*. Here, we explored the opportunity for pathogen-hopping by evaluating a series of quinoxaline-based anti-schistosomal compounds for activity against *P. falciparum*. We identified compounds with low nanomolar potency against 3D7 and multidrug-resistant strains. Evolution of resistance using a mutator *P. falciparum* line revealed a low propensity for resistance. Only one of the series, compound 22, yielded resistance mutations, including point mutations in a non-essential putative hydrolase *pfqrp1,* as well as copy-number amplification of a phospholipid-translocating ATPase, *pfatp2*, a potential target. Notably, independently generated CRISPR-edited mutants in *pfqrp1* also showed resistance to compound 22 and a related analogue. Moreover, previous lines with *pfatp2* copy-number variations were similarly less susceptible to challenge with the new compounds. Finally, we examined whether the predicted hydrolase activity of PfQRP1 underlies its mechanism of resistance, showing that both mutation of the putative catalytic triad and a more severe loss of function mutation elicited resistance. Collectively, we describe a compound series with potent activity against two important pathogens and their potential target in *P. falciparum*.

## INTRODUCTION

Although significant progress has been made in malaria elimination, there were an estimated 249 million new cases and 608,000 deaths due to malaria infection in 2022 [1]. Artemisinin remains the gold standard treatment for uncomplicated malaria, with artemisinin-based combination therapies the dominant treatment since 2005. The emergence of resistance to artemisinin and partner drugs in Southeast Asia, and more recently in Uganda and Rwanda are severe threats to malaria control and elimination [2–4]. New combinations of drugs with novel modes of action can be an effective strategy to delay the emergence of resistance. This requires the identification of new drug targets and the development of new antimalarials, ideally with low propensity for resistance.

To guide antimalarial drug discovery, the Medicines for Malaria Venture (MMV) proposed the type of molecules (Target Candidate Profiles) and medicine (Target Product Profiles) needed [5]. Decades of research and discovery have led to diverse molecules in preclinical and early clinical stages. A common route for antimalarial drug discovery is phenotypic screening, where distinct parasite stages are co-incubated with compounds to identify active molecules. While targets are not necessarily known at this stage in the process, *in vitro* evolution of resistance followed by whole-genome sequencing can be used to deconvolute targets as well as compound mode of action [6].

Quinoxaline derivates are a class of heterocyclic compounds and have been shown to have diverse applications in medicine due to their biological activities. In addition to their known anti-microbial, anti-inflammatory, anti-cancer, anti-depressant, and anti-diabetic activities [7], quinoxaline derivatives also demonstrate anti-plasmodial properties. A Novartis chemical library screen identified BQR695 (2-[[7-(3,4-dimethoxyphenyl)quinoxalin-2-yl]amino]-N-methylacetamide) as an anti-plasmodial compound that acts through inhibition of PfPI4-kinase [8]. In recent work, we showed that quinoxaline compounds possess a low potential for resistance, requiring the use of a mutator parasite line to evolve even low-level resistance [9].

In this study, we explored a pathogen-hopping opportunity, evaluating a series of quinoxaline-containing compounds that have been previously shown to have anti-schistosomal activity [10]. A subset of these compounds have highly potent antimalarial activity, with single-digit nanomolar IC_50_. Using *in vitro* resistance evolution experiments to gain insights into the mechanism of compound action, we identified mutations in quinoxaline resistance protein (PfQRP1), a non-essential putative hydrolase. In addition, we showed that parasites with copy number amplification of a phospholipid-translocating ATPase, *pfatp2*, were also less susceptible to these quinoxaline-based compounds. Overall, we show that quinoxaline-like compounds can be potent anti-infectives, with low propensity for resistance and modest loss of potency in resistant *P. falciparum* parasites. The dual activity against both *Plasmodium* and *Schistosoma* hint at a conserved target or pathway, suggesting further exploration of these compounds in *Plasmodium* may additionally yield insights for accelerating schistosome drug discovery.

## RESULTS

### Activity of quinoxaline compounds against *Plasmodium falciparum*

We investigated the anti-plasmodial activity of a lead anti-schistosomal molecule, compound **22** [10] (**Figure 1A**). This compound showed potent activity against both 3D7 (IC_50_ = 22 nM) and the multi-drug resistant strain Dd2 (IC_50_ = 32 nM). Modification of the nitro group on the C6 position of the central core to either a *N*-acetyl amide (compound **22c**) or *N*-furan-2-carboxamide (compound **22f**) greatly diminished activity against both *P. falciparum* strains (**Figure 1A**), similar to the effect on anti-schistosome activity [10].

**Figure 1:**
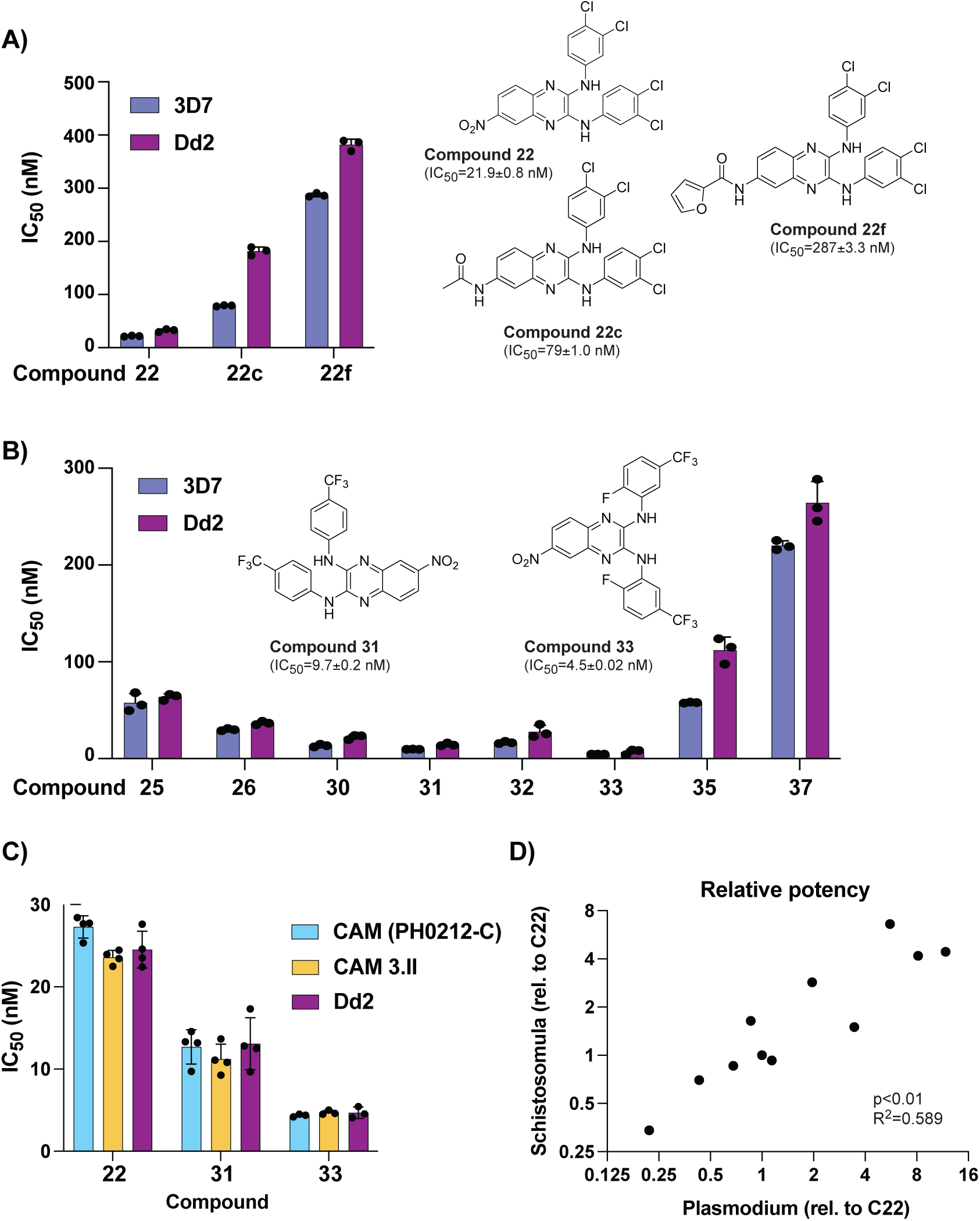
Anti-schistosomal compounds show potent anti-malarial activity. **A)** The lead compound **22** and two derivatives show a range of activities against the *P. falciparum* strains 3D7 and Dd2. Structures of all compounds are in Supplementary Fig. 1, and selected structures are shown with 3D7 IC_50_ values. **B)** Evaluation of additional derivatives of compound **22** identified compounds **31** and **33** as the most potent. **C)** Anti-plasmodial activity of compounds **22**, **31**, and **33** is comparable against multidrug-resistant Cambodian isolates. **D)** Comparison of potency, relative to IC_50_ of compound **22** (referred to as **C22**), between the larva stage of *S. mansoni* (data derived from [10]) and *P. falciparum* 3D7 strain. Raw values for all compounds shown in **Supplementary Table 1**. For (A-C) each dot represents a biological replicate (n=3-4) with mean±SD values shown as a bar chart.

Keeping the C6 nitro group constant, we further explored 8 additional derivatives with modifications of the aromatic rings and the *N*-linker between the quinoxaline core and each aromatic ring (**Supplementary Figure 1** and **Supplementary Table 1**). Introduction of a *N*-ethyl linker (compound **37**) resulted in a ten-fold loss in potency (**Figure 1B**). In contrast, modifications on the aromatic rings strongly influenced phenotypic activity. In fact, the introduction of a trifluoromethyl group in compounds **30-33** led to an increase in potency, with single-digit nanomolar IC_50_ for compounds **31** and **33** (**Figure 1B** and **Supplementary Table 2**). Further evaluation of compounds **22**, **31** and **33** against two Cambodian isolates with multi-drug resistance (to artemisinin, chloroquine, and pyrimethamine) showed no loss of potency relative to the lab strain Dd2 (**Figure 1C**).

Overall, comparison of the activity between *P. falciparum* asexual blood stage parasites and *S. mansoni* schistosomula showed a good correlation across all 11 compounds tested (**Figure 1D** and **Supplementary Table 2**). Based on potencies, three compounds (**22**, **31**, **33**) were subsequently shortlisted to explore mode of action.

### Resistance generation using a mutator parasite

*In vitro* evolution of resistance is a powerful tool for understanding drug mode of action [6]. To facilitate the isolation of mutants, we recently reported a mutator *P. falciparum* line that has an elevated mutation rate and higher propensity to select for resistance, resulting from defective proof-reading due to an introduced mutation in the DNA polymerase δ subunit [9]. To investigate the mode of action of these quinoxaline compounds, the Dd2-Polδ mutator line was pressured with compounds **22**, **31** and **33** (**Figure 2A**). A single-step *in vitro* resistance method was used where triplicate flasks with 1×10^8^ parasites were exposed to 5×IC_50_ of each compound (**Figure 2A**). After 8-10 days of treatment, parasites were undetectable (<0.1% parasitemia) by microscopy for all three compounds. Drug pressure was then removed, and parasites were allowed to recover. After approximately 3 weeks, compound **22**- and **31**-treated parasites recovered in two of the triplicate flasks tested. In contrast compound **33**-treated parasites did not recover even after 60 days.

**Figure 2:**
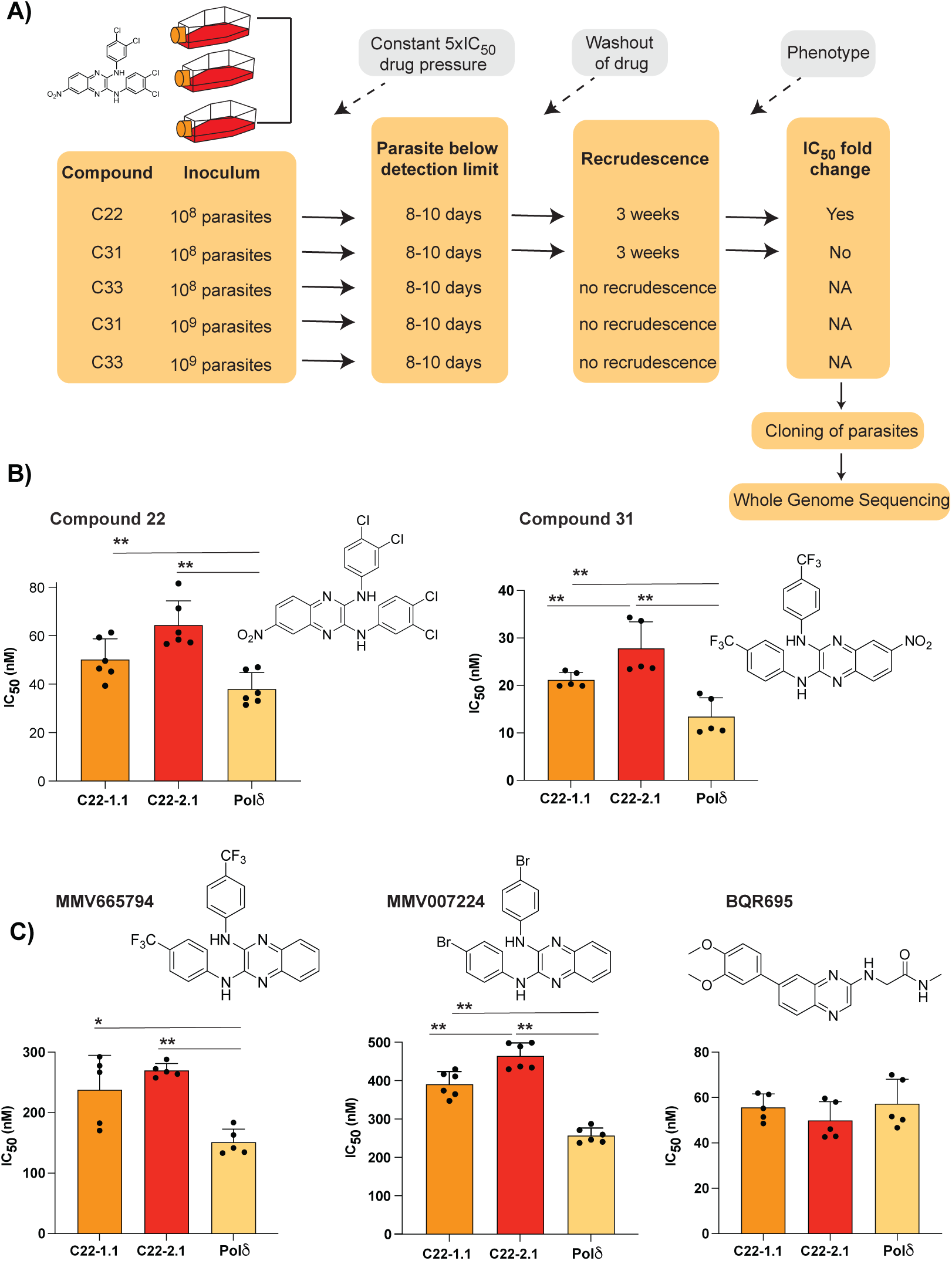
Evolution of resistance to quinoxaline-containing compounds. **A)** Selection scheme showing attempts to generate resistance using the Dd2-Pol*δ* mutator line. Indicated are the compound, parasite inoculum, selection period during which parasites were exposed to 5xIC_50_, recrudescence status of cultures, and shift in IC_50_ of the bulk culture. **B)** Clones from two independent flasks (1.1 and 2.1) from the compound **22** selection were evaluated against compounds **22** and **31**. The parental line, Dd2-Pol*δ* (Pol*δ*) was used as the control. **C)** Clones (1.1 and 2.1) from the compound **22** selection were evaluated against other quinoxaline-containing compounds, MMV665794, MMV007224 and BQR695. Each dot represents a biological replicate (n=4-6) with mean±SD shown as bar chart, and statistical significance determined by Mann-Whitney *U* test (*p<0.05, **p<0.01).

Clonal lines were isolated using limiting dilution of bulk cultures from parasites treated with compounds **22** and **31**. Clones derived from compound-**22** selections from flask 1 (C22-1.1) and flask 2 (C22-2.1) showed a small but consistent shift in IC_50_ to both compound **22** and **31** (**Figure 2B**). In contrast, cultures pressured with compound **31** showed no significant shift in IC_50_ compared to the parental line. To explore if we could generate resistance to compounds **31** or **33**, we repeated the resistance selections using 1×10^9^ parasites as the initial inoculum, however, no recrudescent parasites were recovered (**Figure 2A**). Ramping selections where parasites were cultured initially at 1×IC_50_ and slowly adapted to increasing concentration of drug also yielded no resistance, with parasites unable to proliferate at 2xIC_50_ despite exposing them for almost a month. Thus, the resistance risk with these quinoxaline-based analogues was low, with compounds **31** and **33** proving to be resistance-refractory to date and compound **22** yielding only low-level resistance.

The low propensity for resistance of these quinoxaline compounds was reminiscent of our experience with *in vitro* evolution experiments with two related quinoxaline scaffolds (2,3-dianilinoquinoxaline derivatives without nitro group on the C6 position): MMV665794 (2-N,3-N-bis[3-(trifluoromethyl)phenyl]quinoxaline-2,3-diamine) and MMV007224 (2-*N*,3-*N*-bis(4-bromophenyl)quinoxaline-2,3-diamine) [9, 11]. Examination of cross-resistance of the compound **22**-selected clones to these two compounds revealed a similar low-level shift in IC_50_, suggesting a shared mechanism (**Figure 2C**). In contrast, no cross resistance was found against the PfPI4K inhibitor BQR695 [8], which possesses a quinoxaline core but not the 2,3-dianilino substitution pattern (**Figure 2C**).

### Mutations in PfQRP1 confer resistance

To identify mutations in the compound **22**-selected parasites, we performed whole genome sequencing of eight clones isolated from two separate flasks (clones 1.1-1.4 and clones 2.1-2.4) as well as the sensitive isogenic parent. Only one gene, PF3D7_1359900, was common to all clones, with seven clones encoding a R676I mutation and another encoding a V673D mutation (**Figure 3A** and **Supplementary Table 3**). Notably, this gene was also mutated in previous selections with the compound MMV665794 described above, and encodes a protein we recently designated PfQRP1, for quinoxaline resistance protein [9].

**Figure 3:**
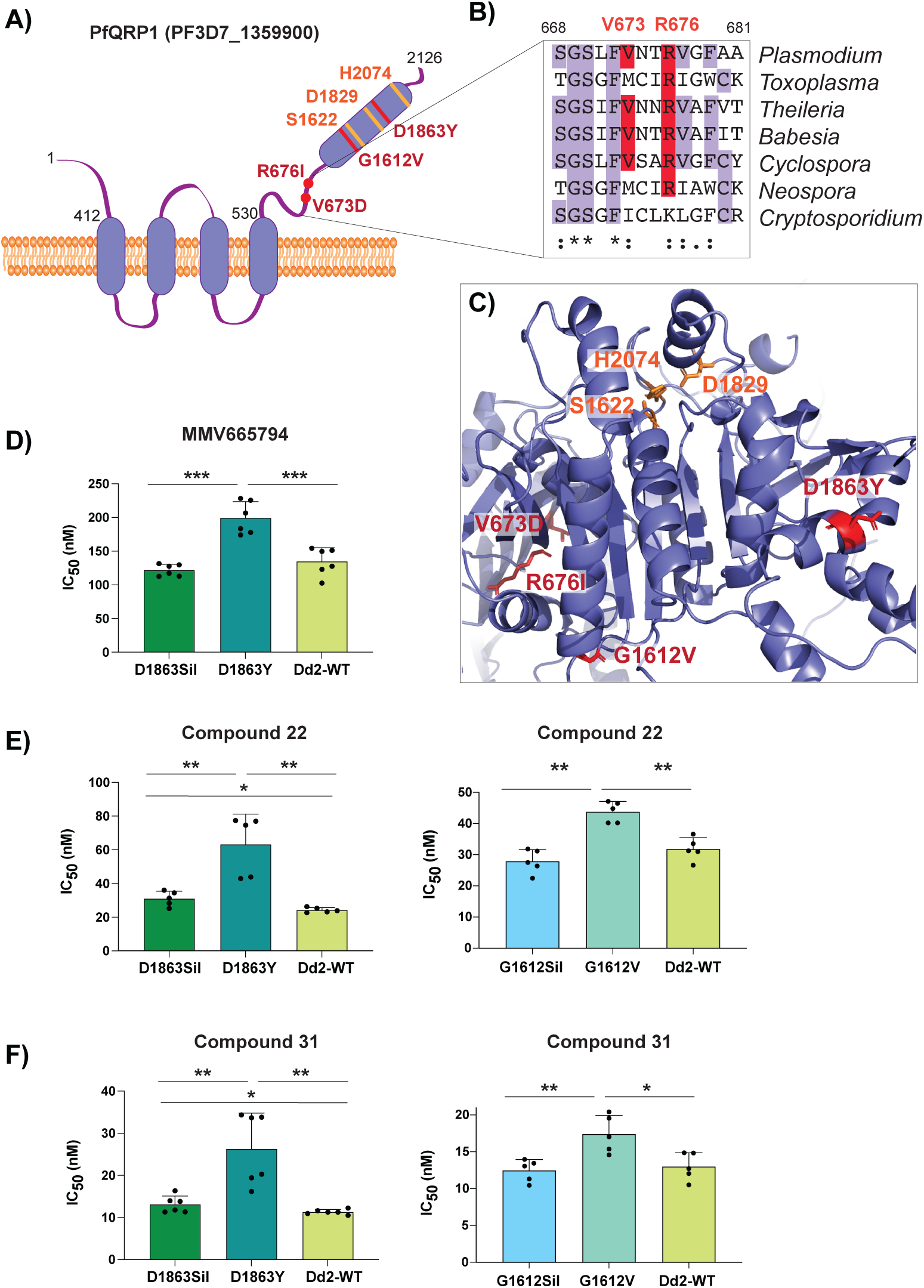
Mutations in PfQRP1 confer resistance to quinoxaline-containing compounds. **A)** Model of the PfQRP1 protein, with 4 predicted transmembrane segments and a putative α/β hydrolase domain. The compound **22**-resistance mutations V673D and R676I, and the MMV665794-resistance mutations G1612V and D1863Y are shown in red, and putative catalytic triad in yellow. **B)** Partial sequence alignment of *pfqrp1* homologs showing partial conservation of residues mutated in drug-selected parasites (red). *P. falciparum pfqrp1* (PF3D7_1359900) with *Toxoplasma gondii* (TGME49_289880), *Theileria parva* (TpMuguga_02g02080), *Babesia bovis* (BBOV_II004840), *Neospora caninum* (NCLIV_042110), *Cyclospora cayetanensis* (cyc_01400) and *Cryptosporidium parvum* (cgd3_590). **C)** AlphaFold model of PfQRP1 showing the resistance mutations (red) and the residues of the putative catalytic triad (yellow). **D-F)** CRISPR-edited parasite lines with the MMV665794-resistance mutations D1863Y and G1612V show a shift against MMV665794 (**D**) as well as compounds **22** (**E**) and **31** (**F**). Control lines with only silent mutations (Sil) or the unedited wild type (WT) are shown. Each dot represents a biological replicate (n=4-6) with mean±SD shown as bar chart, and statistical significance determined by Mann-Whitney *U* tests (*p<0.05, **p<0.01).

Although the function of PfQRP1 is unknown, the 2126 amino acid protein is predicted to encode four transmembrane domains spanning residues 412-530, and a putative alpha-beta hydrolase domain at the C-terminus (**Figure 3A**). The sites of the compound **22**-resistance mutations, V673D and R676I, are relatively well conserved across orthologous Apicomplexan proteins (**Figure 3B**). When mapped onto an AlphaFold-generated structure of PfQRP1, these residues as well as the two previously identified mutations G1612V and D1863Y that confer resistance to MMV665794 [9], are located close to the putative catalytic triad of the hydrolase domain (**Figure 3C**).

To validate the importance of PfQRP1 for resistance to the anti-schistosomal compounds **22** and **31**, we next used previously generated CRISPR-edited mutant lines bearing the G1612V and D1863Y mutations, as well as control edited lines with only silent mutations at those sites. Notably, both compound **22** and **31** showed reduced susceptibility in both CRISPR mutant lines (**Figure 3D-F**).

### Amplification of a lipid flippase confers resistance to quinoxaline compounds

Our observations above demonstrate the importance of PfQRP1 as a resistance mechanism to these quinoxaline-based compounds. However, PfQRP1 is unlikely to be the target, as the gene is non-essential based on mutagenesis in the *piggyBac* screen, as well as the presence of a frameshift mutation near the start of the coding region in one of the MMV007224-resistant clones [9, 12]. Notably, among a small number of copy number variations (CNVs) identified in the compound **22**-resistant lines, clones isolated from flask 2 possessed a CNV of the phospholipid-translocating “flippase” PfATP2 (PF3D7_1219600; **Figure 4A** and **Supplementary Table 4**). Consistent with this, clone 2.1 isolated from flask two had a modestly higher IC_50_ for compound **22** compared to flask one (clone 1.1), despite sharing the same PfQRP1 R676I mutation (**Figure 2B, C**).

**Figure 4:**
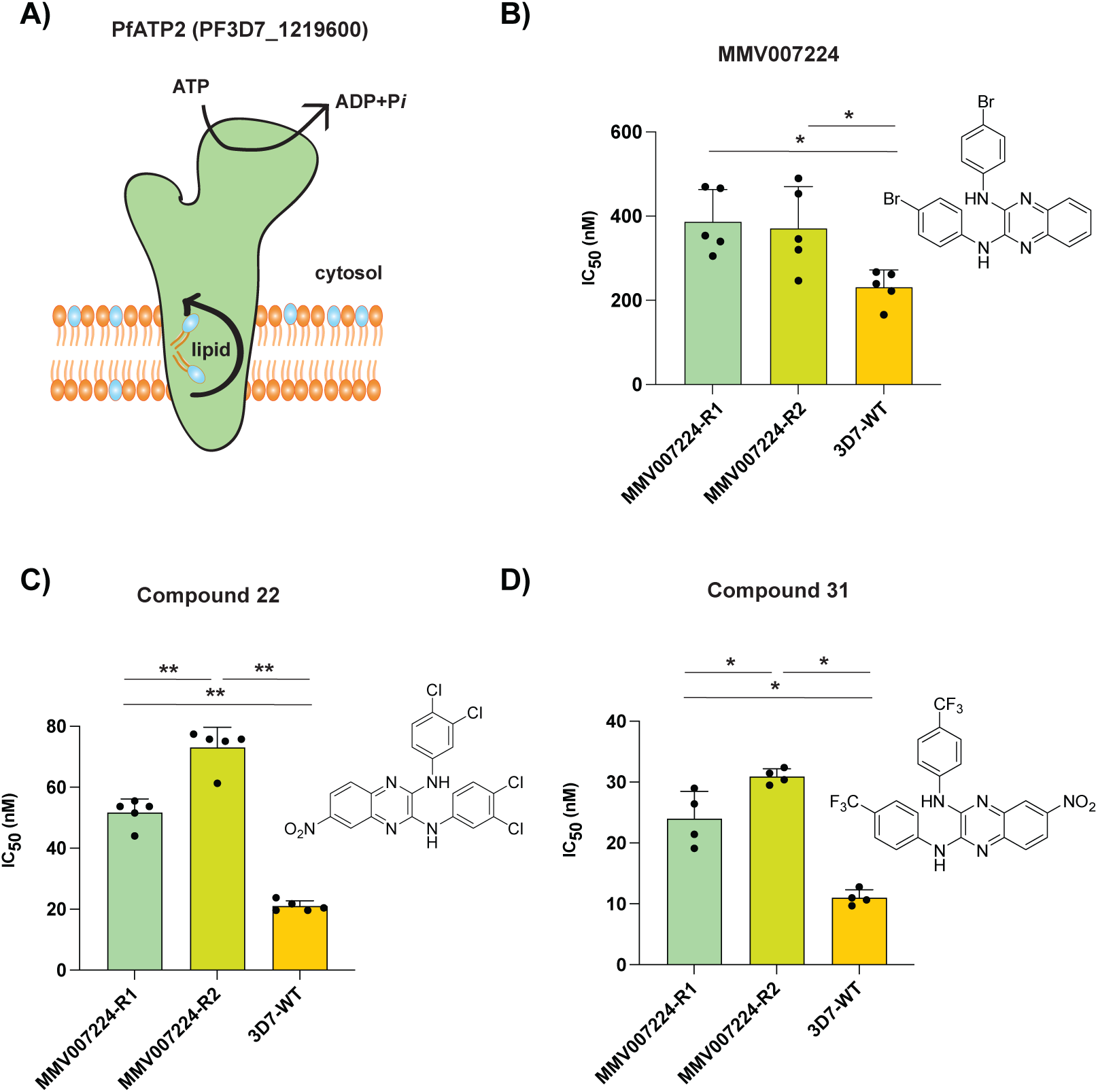
Amplification of a phospholipid-translocating ATPase reduces susceptibility to quinoxaline-containing compounds. **A)** Model of PfATP2 lipid flippase. The P4-ATPase is predicted to contain 10 transmembrane segments, with phospholipid translocation from the luminal to cytosolic leaflet of the membrane powered by ATP hydrolysis. **B-D)** MMV007224-selected clones R1 (*pfatp2* CNV) and R2 (*pfatp2* CNV and *pfqrp1* frameshift) were tested against (**B**) MMV007224, (**C**) compound **22**, and (**D**) compound **31**. Each dot represents a biological replicate (n=4-5) with mean±SD shown as bar chart, and statistical significance determined by Mann-Whitney *U* tests (*p<0.05, **p<0.01).

An association of PfATP2 with related quinoxaline-containing compounds was observed with previous resistance selection experiments with MMV007224. Three independent selections (R1-R3) with MMV007224 all yielded copy number variations (CNVs) covering the *pfatp2* gene, with only R2 also containing an additional *pfqrp1* frameshift mutation at amino acid 100 out of 2126, resulting in a truncated protein [11]. To evaluate whether *pfatp2* amplification also confers resistance to compounds **22** and **31**, we tested the MMV007224-resistant clones R1 and R2. Both lines conferred a 2- to 3-fold increase in IC_50_ for both compounds **22** and **31**, similar to the original selection compound MMV007224 (**Figures 4B-D**). The modestly higher IC_50_ values for line R2 that possesses both the *pfatp2* CNV and the *pfqrp1* frameshift mutation suggest that both mechanisms together may contribute to resistance to quinoxaline-based compounds.

### Loss-of-function mutations in PfQRP1 confer resistance

Finally, we explored whether loss-of-function of the predicted hydrolase activity of PfQRP1 underlies resistance to these quinoxaline-based compounds. Residues corresponding to a potential catalytic triad – S1622/D1829/H2047 in *P. falciparum* – are highly conserved across Apicomplexan orthologs of PfQRP1 (**Figure 5A**) and are in close proximity in the AlphaFold model (**Figure 5B**). We generated a CRISPR-edited truncation mutant at the D1829 residue, inserting a stop codon. Truncation of PfQRP1 within the hydrolase domain resulted in an elevated IC_50_ against compounds **22** and **31**, as well as MMV665794 and MMV007224 (**Figure 5C, D**). Similarly, a more subtle D1829A point mutant behaved similarly to the truncation mutant (**Figure 5C, D**). These results suggest that loss of function in the PfQRP1 hydrolase domain is protective against this compound series.

**Figure 5:**
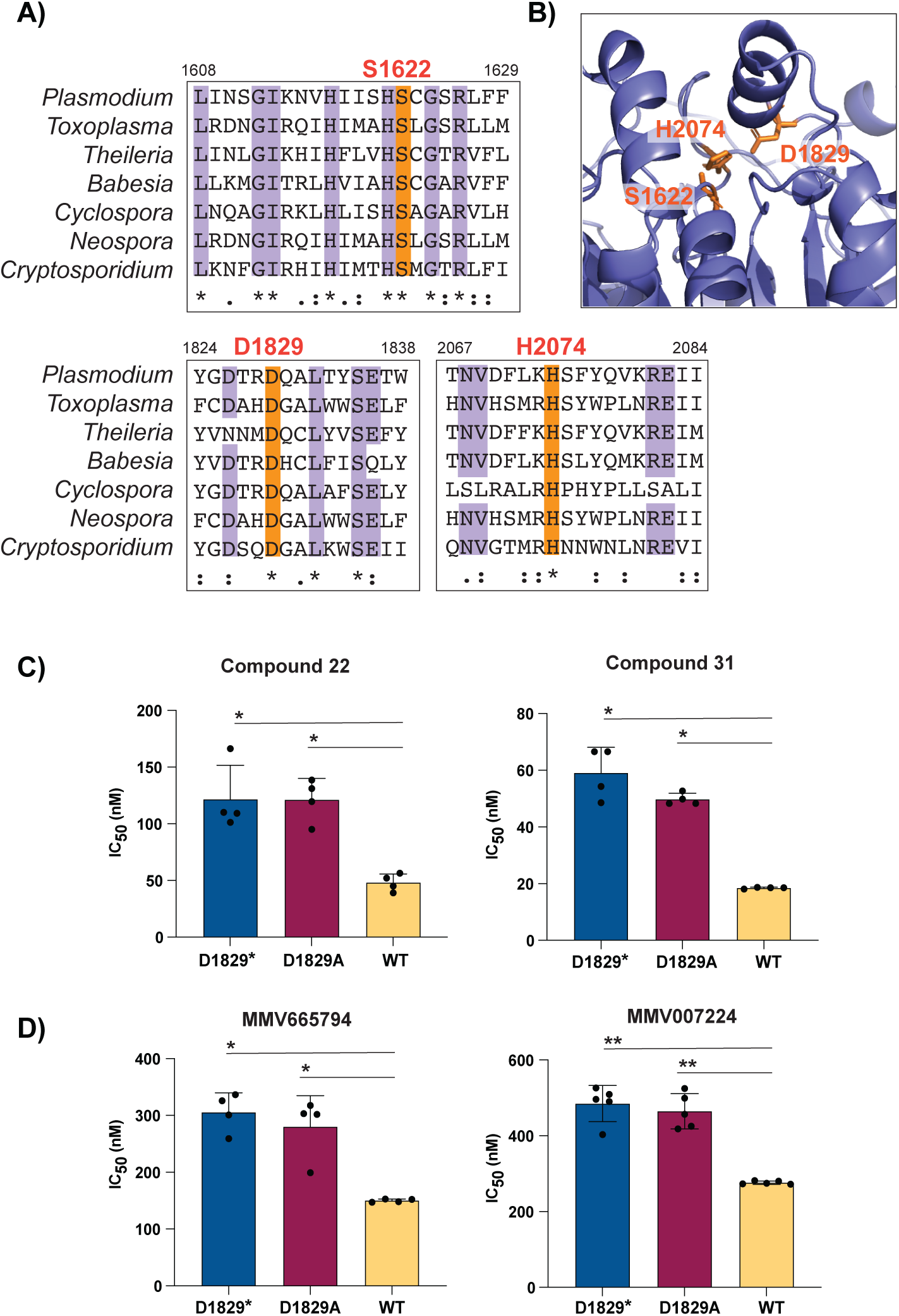
Mutation of the putative catalytic triad of PfQRP1 confers resistance. **A)** Partial sequence alignment of *pfqrp1* homologs showing high conservation of putative catalytic triad residues (orange), comparing *P. falciparum pfqrp1* (PF3D7_1359900) with Apicomplexan orthologs listed in Figure 3. **B)** AlphaFold model of PfQRP1 showing the S-D-H residues that form the putative catalytic triad. **C)** CRISPR-edited parasites with D1829 mutated to a stop codon (D1829*) or alanine (D1829A) show elevated IC_50_ values for **C)** compounds **22** and **31**, and **D)** MMV665794 and MMV007224. Each dot represents a biological replicate (n=4-5) with mean±SD shown as bar chart, and statistical significance determined by Mann-Whitney *U* test (*p<0.05, **p<0.01).

## DISCUSSION

Here, we report that a series of anti-schistosomal, quinoxaline-based compounds have potent activity against the asexual blood stage of *P. falciparum*. The compounds had sub-micromolar antimalarial activity, reaching as low as single-digit nanomolar IC_50_ against lab lines 3D7 and Dd2, as well as multi-drug resistant Cambodian isolates. To understand the mode of action of this compound series, we performed *in vitro* resistance selections with three compounds (**22**, **31**, **33**). We were only successful in generating resistant parasites against compound **22**, and not **31** and **33**, despite using a high inoculum of up to 10^9^ parasites of a mutator line with an elevated mutation rate, which we have shown to elicit resistance to previously irresistible compounds [9]. The inability to generate resistance to compound **31** and **33** using these conditions, and to obtain only low-grade resistance to **22**, suggest a promising resistance profile for this class of compounds and their cognate target.

Whole-genome sequencing of compound **22**-evolved clones identified mutations in *pfqrp1*, a gene we recently characterised as encoding a quinoxaline-resistance protein [9]. We validated PfQRP1 as causal for resistance; CRISPR-edited lines bearing two resistance mutations to another quinoxaline compound, MMV665794 [9] showed cross-resistance to compounds **22** and **31**. PfQRP1 is a large protein of 250 kDa and is predicted to possess four N-terminal transmembrane segments as well as a putative C-terminal alpha/beta hydrolase domain. Although the mechanism of resistance is unknown, the compound **22**-resistance mutations, as well as the two previously identified mutations conferring resistance to MMV665794, map near to the putative hydrolase domain in the AlphaFold structure. In addition, the frameshift mutation identified in the MMV007224-resistant clone R2 suggests that resistance may be mediated by PfQRP1 loss-of-function.

To further explore whether the putative hydrolase function of PfQRP1 was related to its mechanism of resistance, we generated a point mutation in D1829, one of the putative catalytic triad residues. Conversion to alanine, or a more dramatic stop codon insertion, both conferred a similar level of resistance to the drug-selected mutations, suggesting that loss of PfQRP1 hydrolase function mediates protection. Whether this is through direct action on the compounds, as with the PfPARE prodrug convertase [13], or a more indirect mechanism is unknown.

The non-essential nature of PfQRP1 suggests that the direct target of these quinoxaline-based compounds is likely another protein, with the phospholipid-transporting P4-ATPase, PfATP2 being a prime candidate. Copy number amplification of *pfatp2* was observed in a subset of compound **22**-selected clones, suggesting that increased abundance of this putative target is protective. Similarly, *pfatp2* CNVs were previously obtained with resistance selections using MMV007224 [11]. Our data showed that a MMV007224-resistant clone with a *pfatp2* CNV but no PfQRP1 mutation also confers cross-resistance to compounds **22** and **31** described herein. In general, phospholipid-transporting “flippases” perform ATP-dependent translocation of phospholipids from the extracellular or luminal face of the lipid bilayer towards to cytoplasmic leaflet [14–16]. Unlike *pfqrp1*, *pfatp2* is likely an essential gene based on the absence of *piggyBac* insertions and the inability to disrupt the *P. berghei* ortholog, which is localised to the parasite plasma membrane or parasitophorous vacuolar membrane [17, 18]. Furthermore, P-type ATPases are well-established drug targets, with Na^+^-ATPase PfATP4 established as the target of cipargamin, which is in phase 2 clinical trials [19, 20]. The significance of the small number of other CNVs is unknown, although the presence of an epigenetic regulator, *bdp4*, in one CNV might affect expression of genes related to the mode-of-action of these quinoxaline compounds.

The correlation between phenotypic activity against *Plasmodium* asexual blood stages and *Schistosoma* schistosomula for the 11 compounds tested is suggestive of a shared mode of action. In addition, evaluation of the 400 compounds from the Malaria Box against *S. mansoni* identified MMV007224 as among the most active hits against adult worms [21]. Notably, another of the most active compounds against adult *Schistosoma* from the Malaria Box was MMV665852, a N,N′-diarylurea compound. Resistance selections in *P. falciparum* with MMV665852 also yielded CNVs in *pfatp2* [11]. Although we could not identify a clear homolog of PfQRP1, there are multiple phospholipid-transporting ATPases annotated in the *Schistosoma mansoni* genome. InterPro domain IDs (IPR001757 and IPR006539 for a generic P-type ATPase and type 4 ATPase, respectively) identify 20 P-type ATPases in *S. mansoni*, of which 6 belong to the P4-ATPase subfamily that specialise in lipid rather than ion transport (**Supplementary Figure 2**). These may warrant further investigation as a new target class.

In summary, we describe the dual-parasite activity of a quinoxaline-core compound series that is potent against both *P. falciparum* and *S. mansoni*. *In vitro* evolution of resistance using a *P. falciparum* mutator line indicates this series has a low propensity for resistance, and when achieved, only a modest loss of potency. Resistance can be driven by mutations in a quinoxaline-resistance protein, PfQRP1, or CNVs of a phospholipid-transporting ATPase, PfATP2. The correlation in phenotypic activity between *Plasmodium* and *Schistosoma* hint at a shared mode of action and potential new targets for controlling the parasites responsible for two important infectious diseases.

## MATERIALS AND METHODS

### Parasite cultivation and transfection

*P. falciparum* parasites were cultured at 3% hematocrit in RPMI 1640 (Gibco) medium consisting of 0.5% Albumax II (Gibco), 25mM HEPES (Sigma), 1x GlutaMAX (Gibco), 25μg/mL gentamicin (Gibco). A tri-gas mixture (1% O_2_, 3% CO_2_, and 96% N_2_) was used for parasite cultures. Parasites were cultured in fresh human erythrocytes obtained with ethical approval from anonymous healthy donors from the National Health Services Blood and Transplant (NHSBT) or the Scottish National Blood Transfusion Service (SNBTS). Parasites were synchronized using a sorbitol and Percoll density gradient method [22]. Transfection was performed using ring-stage parasites (5-8% parasitemia) using a Gene Pulser Xcell (Biorad) electroporator.

To generate the PfQRP1 hydrolase mutants (D1829stop and D1829A), the pDC2-Cas9-gRNA-donor plasmid [23] was transfected into the *P. falciparum* Dd2 line. Donor plasmid (50 µg) containing the sgRNA (AGCTTTAACATATTCAGAAA) and a *pfqrp1* homology region of 672 bp, centred on the D1829 residue, was used for transfection. Transfectants were selected using 5 nM WR99210 (a potent inhibitor of *P. falciparum* dihydrofolate reductase (DHFR) provided by Jacobus Pharmaceuticals) for 8 days, followed by no drug treatment until parasites were observed. Parasite clones were obtained using limiting dilution cloning. Confirmation of gene editing was performed with allele-specific PCR (forward primer CTGAAGAAGATGAATGGGAACA and reverse primer CCACCTTCTCCTTCACCAAC) and Sanger sequencing using an internal primer (TGGAAGAAAGGAAAACACAACA).

### *In vitro* drug resistance selections using Dd2-Polδ

*In vitro* evolution of resistance was performed against compounds **22**, **31** and **33**. The mutator parasite line Dd2-Polδ was used for resistance generation [9]. Three independent flasks with either 1×10^8^ or 1×10^9^ parasites each were treated with 5×IC_50_, and parasite death was monitored by microscopy (Giemsa staining). Viable parasites were undetectable by blood smear after eight days of compound treatment. After eight days, compounds were removed from the media and the media was changed on alternate days. Parasites reappeared after approximately 3 weeks from the washout of the drug (**Figure 2A**). For recrudescent cultures, compound susceptibility was determined by dose-response assays. Parasite clones were obtained by limiting dilution and harvested for genomic DNA extraction, followed by whole genome sequencing.

### Compound dose-response assay

Compound dose-response assays were performed in flat-bottom 96-well plates. Dose-response assays were performed with strains 3D7, Dd2, CAM (PH0212-C/K13^C580Y^; [24]) and Cam3.II (PH0306-C/K13^R539T^; [25]). Tightly synchronized ring-stage parasites were diluted to 1% parasitemia at 2% hematocrit, and incubated with a two-fold serial dilution of compounds in complete medium. Untreated parasites and red blood cells (RBCs) only were included in the assay plate as controls. Parasite growth was assessed after 72 hours by lysing parasites using 2× lysis buffer consisting of 10 mM Tris-HCl, 5 mM EDTA, 0.1% w/v saponin, and 1% v/v Triton X-100, supplemented with 2× SYBR Green I (Molecular Probes). The fluorescence was measured using a FluorStar Omega v5.11 plate reader. IC_50_ analysis was performed using GraphPad Prism v9, and statistical significance was determined by a two-sided Mann-Whitney *U* test. All assays were performed in technical duplicate with at least three biological replicates, as noted in the figure legends.

### Whole genome sequencing

Parasites were harvested using 0.1% saponin lysis buffer of RBCs, followed by three washes of PBS. Genomic DNA was extracted using the DNAeasy Blood and Tissue Kit (Qiagen). Genomic DNA concentration was quantified using a Qubit dsDNA BR assay kit and measured using a Qubit 2.0 fluorometer (Thermo Fisher Scientific).

### Single nucleotide variant and copy number variant calling

Whole-genome sequencing was performed using a IDT-ILMN Nextera DNA UD library kit and multiplexed on a NextSeq flow cell to generate 150 bp paired-end reads. Sequences were aligned to the *Pf* 3D7 reference genome (PlasmoDB-48; https://plasmodb.org/plasmo/app/downloads/release-48/Pfalciparum3D7/fasta/) using the Burrow-Wheeler Alignment (BWA version 0.7.17). PCR duplicates and unmapped reads were filtered out using Samtools (version 1.13) and Picard MarkDuplicates (GATK version 4.2.2). Base quality scores were recalibrated using GATK BaseRecalibrator (GATK version 4.2.2). GATK HaplotypeCaller (GATK version 4.2.2) was used to identify all possible single nucleotide variants (SNVs), filtered based on quality scores (variant quality as function of depth QD > 1.5, mapping quality > 40, min base quality score > 18, read depth > 5) and annotated using SnpEff version 4.3t [26]. Comparative SNP analyses between eight drug-treated Dd2-Polδ clones and the Dd2-Polδ parental strain were performed to generate the final list of SNPs (**Supplementary Table 3**). BIC-Seq version 1.1.2 [27] was used to discover copy number variants (CNVs) against the Dd2-Polδ parental strain using the Bayesian statistical model. SNPs and CNVs were visually inspected and verified using Integrative Genome Viewer (IGV). All gene annotations in the analysis were based on PlasmoDB-48 (https://plasmodb.org/plasmo/app/downloads/release-48/Pfalciparum3D7/gff/).

## Data availability

All associated sequence data are available at the European Nucleotide Archive under accession code PRJEB74174.

## Acknowledgements

We would like to thank R. Fairhurst for providing the Cambodian strains. This work was supported by funding to MCSL from Wellcome (206194/Z/17/Z). D.A.F. gratefully acknowledges support from the NIH (R01 AI124678). KFH and AB acknowledges support from the Life Sciences Wales Research Network, a Welsh Government Ser Cymru initiative.

## Author contributions

MR, GP, KFH and MCSL conceived the study. MR performed the *in vitro* drug selection experiments and parasite transfections. MR and MCSL performed the drug sensitivity assays. Whole-genome sequencing and data analysis were performed by TY and DAF. MR, GP, KFH and MCSL planned the experiments and DAF, AB, KFH and MCSL supervised the study. All authors contributed to writing the paper.

---

**Supplementary Figure 1:**
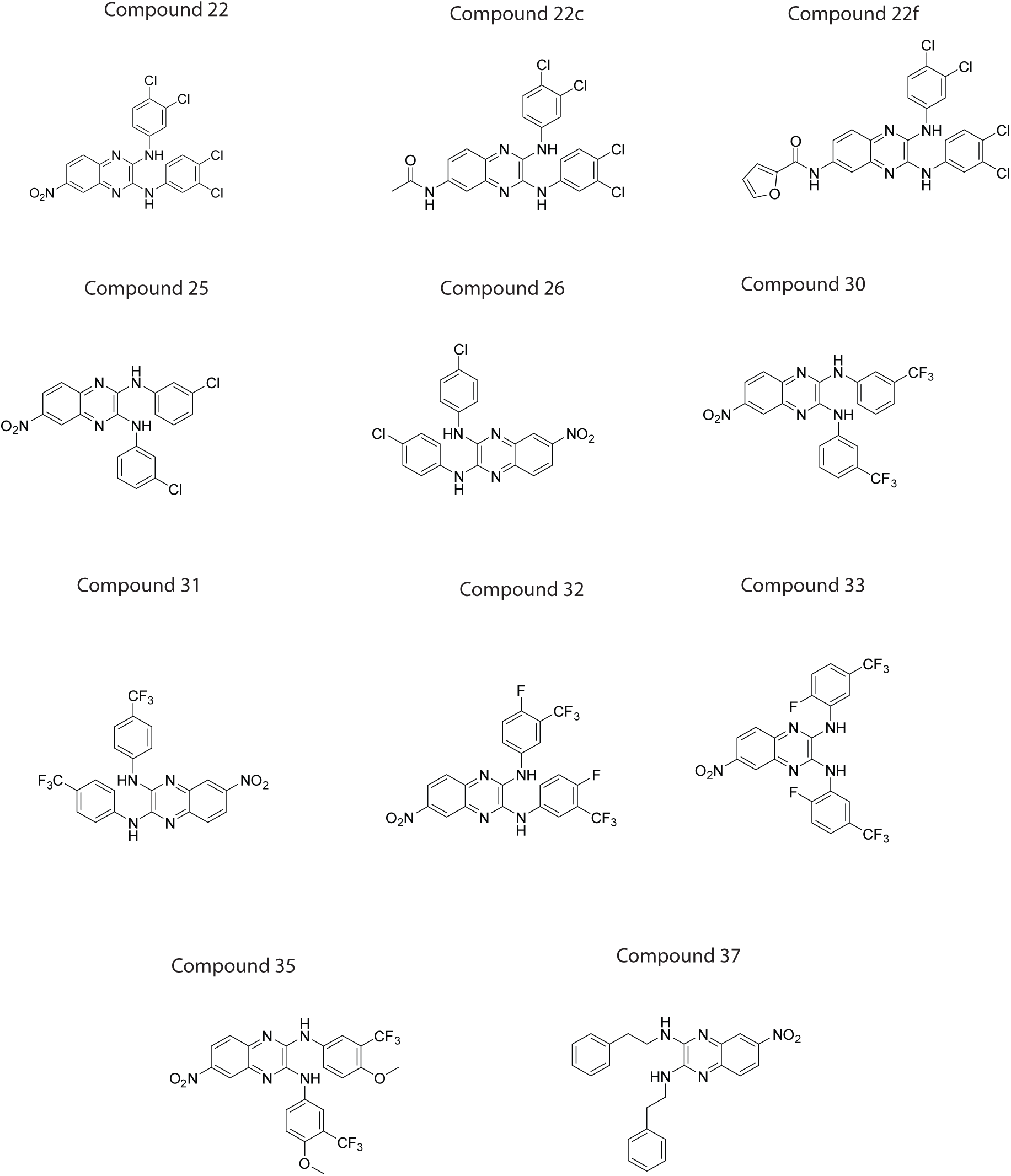
Structures of compounds.

**Supplementary Figure 2:**
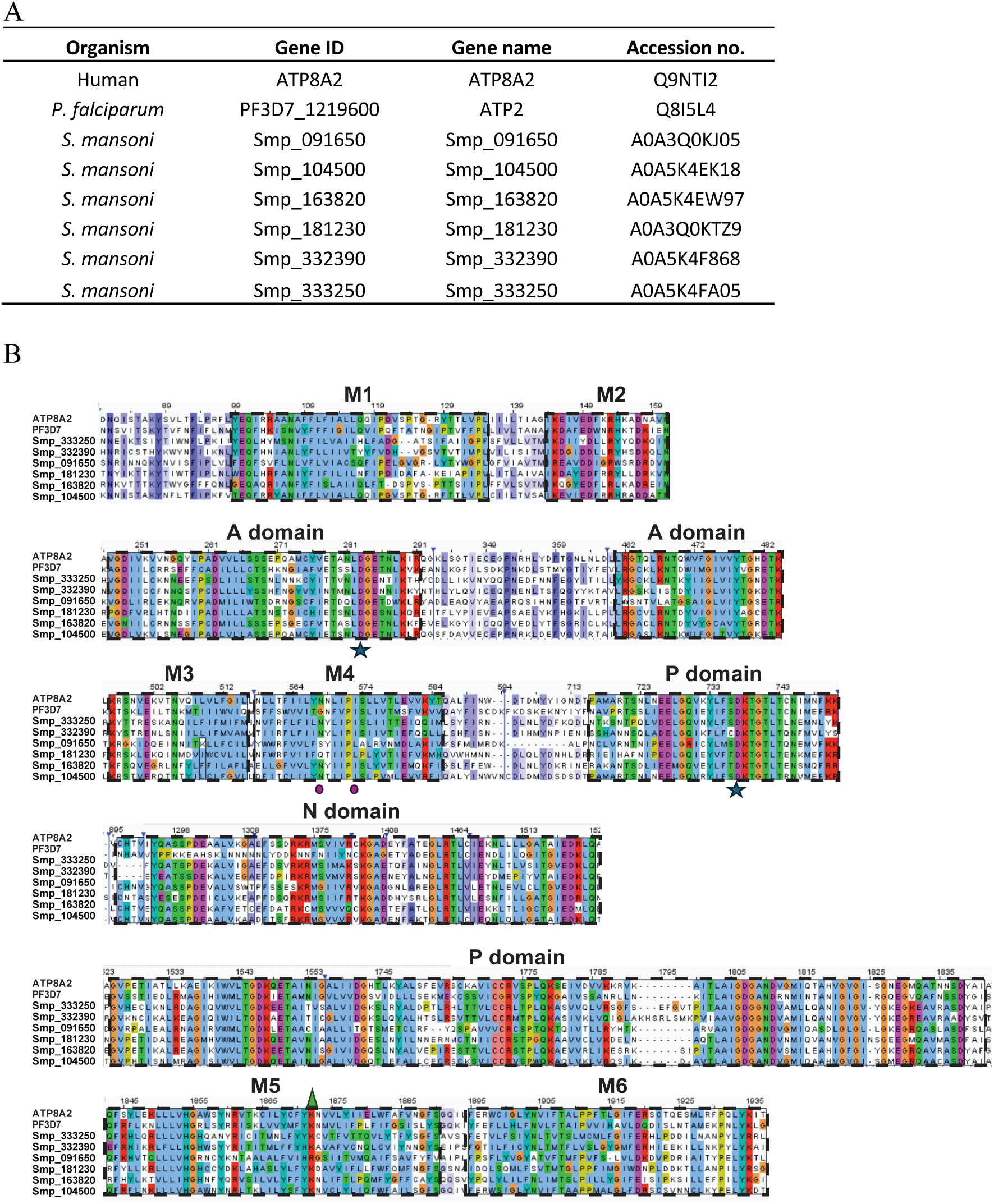
Sequence alignment of phospholipid flippases. **A)** Lipid flippases used in the alignment. **B)** Sequence alignment of human ATP8A2, *P. falciparum* PfATP2, and six putative lipid-translocating ATPases from *S. mansoni*. The actuator (A), nucleotide binding (N), and phosphorylation (P) domains are shown, as well as the first six transmembrane segments (M1-6). Key conserved residues D (in A domain) and E (P domain) involved in the phosphorylation (DKTGT) and dephosphorylation (DGET) cycle are highlighted by a star. The purple circles highlight the conserved N and I residues located in M4 domain that are important for recognition and release of lipid, respectively. The green triangle indicates the K residues in the M5 domain required for the sensitivity to the lipid subtype [14, 28].

## List of Supplementary Tables

**Supplementary Table 1:** Compound information.

**Supplementary Table 2:** Activity against *Plasmodium* asexual blood stages, *Schistosoma* schistosomula and HepG2 cells.

**Supplementary Table 3:** Single-nucleotide variants for compound **22**-selected clones.

**Supplementary Table 4**: Copy number variants from compound **22**-selected clones.

